# Long-term single-particle tracking by NIR imaging using Au_42_ (gold) quantum needles

**DOI:** 10.64898/2026.06.24.734378

**Authors:** Shunsuke Yagi, Shinjiro Takano, Ryo Nishiyama, Ryosuke Oketani, Tatsuya Tsukuda, Kotaro Hiramatsu

## Abstract

Single-particle tracking (SPT) over time enables direct observation of molecular transport and interactions in living cells. Fluorescence-based SPT has provided insights into intracellular processes such as endocytosis, receptor signaling, and drug delivery. Extending the observation window to several hours and beyond is critical for capturing slow intracellular dynamics, including the full course of endosomal trafficking, the long-term accumulation of particles within subcellular compartments, and transitions between transport modes that occur on hour-scale timescales. However, long-term intracellular SPT under visible-wavelength excitation remains challenging because fluorescence probes generally suffer from photobleaching and phototoxicity. While near-infrared (NIR) excitation can simultaneously mitigate these issues, generally weak emission of NIR-emitting dyes has hindered its wide application in long-term SPT. Here, we demonstrate long-term NIR SPT using atomically precise gold quantum needles, Au_42_(PET)_32_ (PET = 2-phenylethanethiolate). Continuous tracking of intracellular particles in living HEK293 cells was achieved for up to 12 h. Trajectory analysis revealed temporal transitions between directional and diffusive transport, as well as the accumulation of multiple particles within localized intracellular domains over several-hour timescales. The high photostability of Au_42_, combined with low phototoxicity of NIR excitation, enables visualization of intracellular transport dynamics over timescales difficult to access using conventional visible fluorescent probes. These results establish Au_42_-based NIR imaging as a platform for long-term, low-phototoxicity intracellular SPT and provide a framework for investigating slow intracellular dynamics in living systems.

## 1. Introduction

Single-particle tracking (SPT) is a powerful approach for investigating molecular transport and interactions in heterogeneous environments such as living cells.^1,2^ SPT follows individual labeled particles over time, providing access to the heterogeneous dynamics of individual particles at high spatial and temporal resolution. Fluorescence-based SPT has played a central role because it offers high sensitivity and enables the simultaneous detection of multiple probes.^3–5^ This capability has provided insights into diverse intracellular processes, including drug delivery,^6^ endocytosis dynamics,^7^ and receptor signaling.^8^ In recent years, SPT techniques capable of tracking individual particles over several hours have been developed.^9^ Long-term SPT is essential for capturing the full course of intracellular processes, such as the trajectory of endosomal trafficking, the long-term accumulation of particles within subcellular compartments, and transitions between distinct transport modes that occur over hour-long timescales.

Despite these capabilities, conventional SPT still faces significant challenges in long-term measurements. One major limitation is phototoxicity, which restricts the duration of live-cell observations during prolonged imaging. The other major limitation is that photobleaching of fluorescent probes reduces signal intensity over time, thereby limiting detection sensitivity. These issues are closely associated with the use of visible-wavelength excitation. In addition, visible excitation induces cellular autofluorescence, which increases background signals and fundamentally limits detection sensitivity. In contrast, NIR excitation can substantially reduce phototoxicity and cellular autofluorescence, making it advantageous for long-term live-cell imaging. Several studies have demonstrated SPT using NIR probes, including single-walled carbon nanotubes (SWCNTs) and upconversion nanoparticles (UCNPs).^7,10–13^ However, no existing probe system simultaneously satisfies the requirements of high brightness, photostability, and low invasiveness for long-term intracellular SPT. Consequently, stable long-term tracking of intracellular particles remains challenging. The development of such probes would broaden the range of slow intracellular transport processes accessible to quantitative long-term SPT.

Recently, highly anisotropic gold nanoclusters have attracted much attention due to their intense NIR absorption/emission and high photostability.^14^ Among them, a series of needle-shaped gold nanoclusters, gold quantum needles (Au QNs) or gold quantum rods (Au QRs), having a pile of Au_3_ triangles in their Au core is the good candidate for the NIR probes.^15–18^ In order to unveil their applicability for long-term SPT, Au_42_(PET)_32_ (PET = 2-phenylethanethiolate) gold nanocluster (Au_42_ in short) was chosen as a benchmark.^19,20^ Au_42_ exhibits both absorption and emission in the NIR region (λ_abs_ = 806 nm; λ_emi_ = 875 and 1040 nm, in dichloromethane (DCM)). These features are attractive for an SPT probe: NIR excitation and emission support low-phototoxicity long-term imaging, while structural precision ensures batch-to-batch reproducibility of the photophysical properties.

In this study, we demonstrate long-term NIR-SPT using Au_42_, enabling continuous tracking of individual particles in living cells for up to 12 h. To introduce Au_42_ into living cells, we developed water-dispersible probes (Adots). We then constructed a fluorescence microscopy system optimized for long-term NIR imaging. Using this system, we continuously tracked individual Adots in live HEK293 cells over 12 h. Trajectory analysis revealed the accumulation of multiple particles within localized intracellular regions, consistent with vesicle-mediated intracellular transport along the endocytic pathway. These results demonstrate that Au_42_ provides a robust platform for long-term, minimally invasive SPT and offers an experimental framework for investigating intracellular transport dynamics over extended periods.

## 2. Results and Discussion

### 2-1. Development and characterization of fluorescent probes based on gold quantum needles

Pristine Au_42_ exhibits limited water solubility and cannot be directly introduced into cell culture media. To improve the water solubility of the probe, Au_42_ clusters were encapsulated into 60 nm polystyrene beads with hydrophilic surface modification to yield water-dispersible Au_42_-based nanoparticles (see Materials and Methods, Section 4-1), hereafter referred to as Adots. Although this approach increases the probe size compared to individual clusters, it enables the introduction of Au_42_ into the cells and provides a defined matrix that protects Au_42_ from direct contact with the cellular environment.

We first examined whether the NIR optical properties of Au_42_ are retained after encapsulation into polystyrene beads by measuring the absorption and emission spectra of Adots (Fig. 1A). The absorption peak of Adots appeared at 800 nm, which is nearly identical to that of Au_42_ in DCM (806 nm).^19^ The emission peaks of Adots appeared at 870 nm and 1080 nm. The 870 nm peak is consistent with that of Au_42_ (875 nm in DCM), whereas the 1080 nm peak is red-shifted compared to Au_42_ (1040 nm in DCM).^19^ Because the shift of the 1080 nm emission peak is small, it is likely attributable to a solvent effect arising from the difference between the polystyrene environment and DCM. These results indicate that encapsulation of Au_42_ into polystyrene beads does not alter its characteristic NIR absorption and emission properties, as previously confirmed by the encapsulation in the bulk polystyrene film.^19,20^ This suggests that the chemical state of Au_42_, including its ligands, is largely preserved inside the beads.

**Figure 1.**
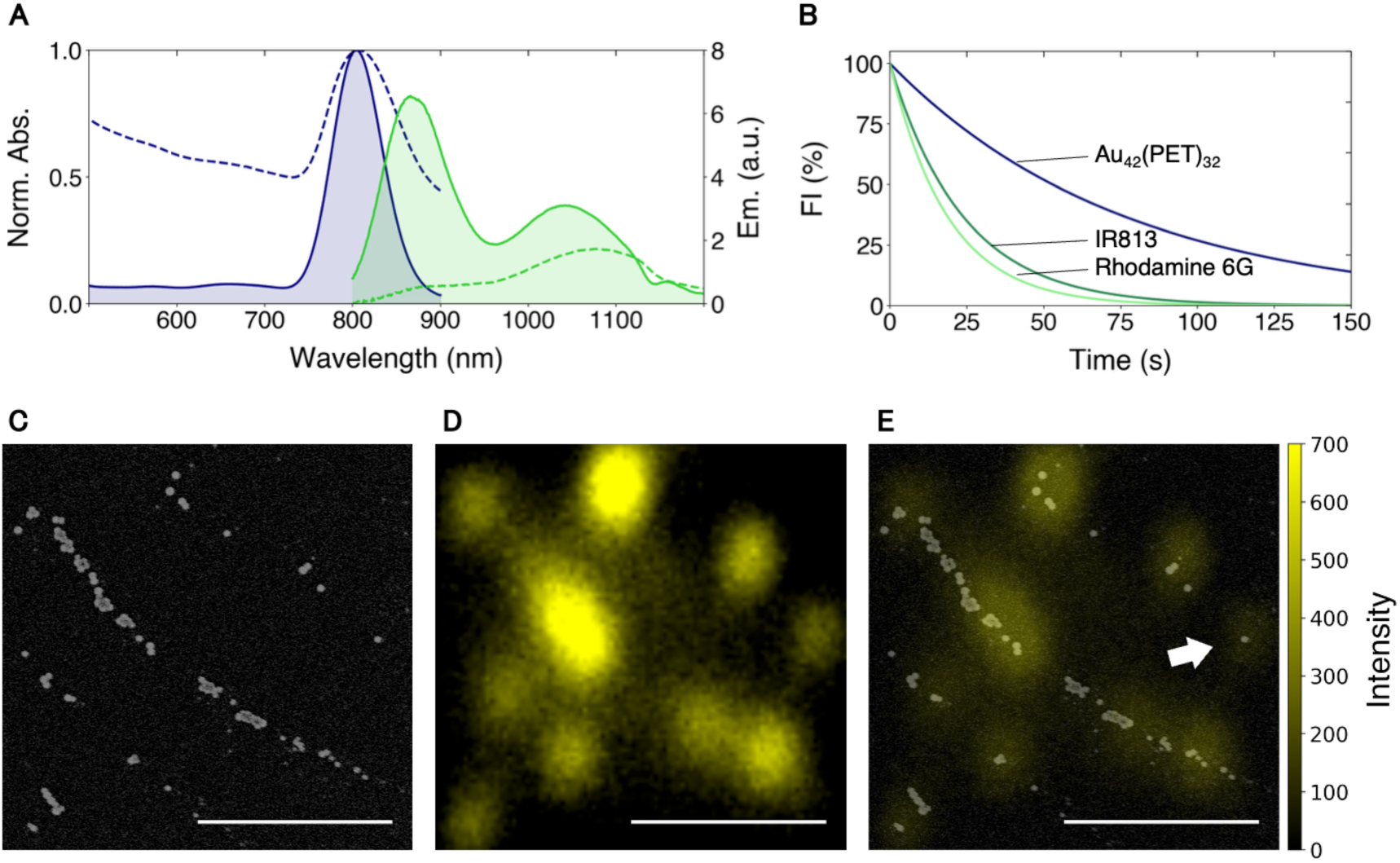
Characterization of Adots (Au_42_(PET)_32_ encapsulated in 60 nm polystyrene beads). (A) Absorption and emission spectra of Au_42_ encapsulated in 60 nm polystyrene beads (dashed lines) and Au_42_ dissolved in DCM (solid lines). The emission spectra were measured under 785 nm excitation. (B) Photobleaching analysis of Au_42_ in comparison with Rhodamine 6G and IR813. Each dye was encapsulated at the same concentration within 60 nm polystyrene beads. The temporal evolution of fluorescence intensity (FI) under continuous-wave excitation is shown. Au_42_ and IR813 were excited at 785 nm, whereas Rhodamine 6G was excited at 532 nm. To match the photon flux between these conditions, irradiation was performed at 0.64 mW for 785 nm and 0.43 mW for 532 nm. (C) SEM image of Os-coated Adots deposited on a glass substrate, acquired at an accelerating voltage of 1.5 kV. (D) Adots fluorescence image of the same region as in (C). (E) Overlay of the SEM image and the corresponding fluorescence image. All scale bars, 2 μm.

Next, the photostability of Au_42_ was assessed by monitoring its fluorescence decay under continuous excitation (Fig. 1B). For comparison, Rhodamine 6G (visible fluorophore) and IR813 (NIR fluorophore) were also encapsulated into 60 nm polystyrene beads, and their fluorescence intensity changes were measured under identical conditions (Fig. 1B). The local concentration of all fluorophores inside the beads was adjusted to 15 mmol/L. Au_42_ and IR813 were excited at 785 nm, while Rhodamine 6G at 532 nm (continuous wave). The laser power measured immediately after the objective lens was 0.64 mW for 785 nm and 0.43 mW for 532 nm. The laser powers were adjusted so that the photon flux was equivalent at the two wavelengths. The single-exponential decay curves were observed in all samples and their decay constants were obtained as 76.3 s (Au_42_), 23.8 s (IR813), and 18.8 s (Rhodamine 6G), respectively. These results indicate that Au_42_ exhibits higher photobleaching resistance than both NIR fluorophores and the visible fluorophore.

The size and morphology of Adots were characterized by scanning electron microscopy (SEM) (Fig. 1C). An aqueous dispersion of Adots was drop-cast onto a glass substrate and allowed to dry prior to measurement. Spherical particles with diameters of approximately 60 nm were observed and are considered to correspond to Adots. The retention of the ∼60 nm size indicates that encapsulation does not significantly perturb the parent polystyrene beads, and the sub-100-nm size lies well within the range efficiently internalized by cells. Non-spherical objects and structures with the sizes of approximately 10 nm were also observed. These features are attributed either to residual Pluronic F-127 surfactant or to Au42-containing micellar aggregates formed by Pluronic F-127.^21^

We finally verified at the single-particle level that the particles observed in the SEM image were fluorescent by acquiring a fluorescence image from the same region shown in Fig. 1C using a home-built fluorescence microscope (Fig. 1D, see Materials and Methods, Section 4-4). The identity of the field of view in the fluorescence and SEM images was confirmed by imaging a marker pattern fabricated on the substrate. The overlay of the two images is shown in Fig. 1E. Fluorescence signals were observed from the spherical particles identified in the SEM image, confirming that these particles correspond to fluorescent Adots. Furthermore, as indicated by the arrows in Fig. 1E, fluorescence from individual particles could be detected. Not all particles exhibited detectable fluorescence, likely due to variations in the amount of encapsulated Au_42_ among particles. One possible cause of this inhomogeneity is that the encapsulation terminates before Au_42_ clusters uniformly distribute throughout all polystyrene beads during the encapsulation process (see Materials and Methods, Section 4-1), resulting in particle-to-particle differences in the local concentration of Au_42_. This issue could potentially be mitigated by repeating the Au_42_ encapsulation procedure to increase the uniformity of probe loading among particles.

### 2-2. Single particle tracking by long-term NIR imaging using Au_42_

To demonstrate long-term SPT using Au_42_, we first performed live-cell imaging for 12 h after Adots were taken up by HEK293 cells by endocytosis (see Materials and Methods, Section 4-3). These cells were imaged using a home-built wide-field microscopy system (see Materials and Methods, Section 4-4). Fluorescence images of Adots with 785-nm excitation and bright-field images of the cells at 940 nm were recorded at 1-min intervals for a total duration of 12 h. To maintain the focal plane over this extended period, an autofocus system was deployed into the microscope (see Materials and Methods, Section 4-4).

Figure 2A shows bright-field and fluorescence images of HEK293 cells that have taken up Adots. In the fluorescence images, punctate fluorescence signals are detected in each frame. The variations in spot size and intensity presumably originate from differences in the number of Au_42_ clusters in each Adot particle, which may arise from inhomogeneous environments during Adot synthesis and from aggregation of Adots within the cellular environment. The fluorescence background levels at the extracellular medium (Point 1 in Figs. 2B, 2C) and the cell body (Point 3 in Figs. 2B, 2C) were negligible, whereas fluorescence signals from intracellular Adots (Points 2 and 4 in Figs. 2B, 2C) were significantly higher than the background. These results indicate that cellular autofluorescence under NIR excitation is negligible compared with the fluorescence intensity of Adots, thereby enabling high-contrast imaging of intracellular particles. In addition, no apparent cell death or detachment was observed during the 12-h measurement under 785-nm excitation, suggesting that phototoxic effects under the present imaging conditions were sufficiently low to maintain the cell viability. Moreover, after the 12-h imaging, several Adots fluorescence spots remained distinguishable from the background. The signal-to-noise ratios (SNRs) were 15.6 and 19.7 for particles 1 and 2, respectively (Figs. 2D, 2E). These results demonstrate that the combination of NIR excitation, the high brightness of Adots, and the strong photobleaching resistance of Au_42_ enables long-term continuous imaging in live cells.

**Figure 2.**
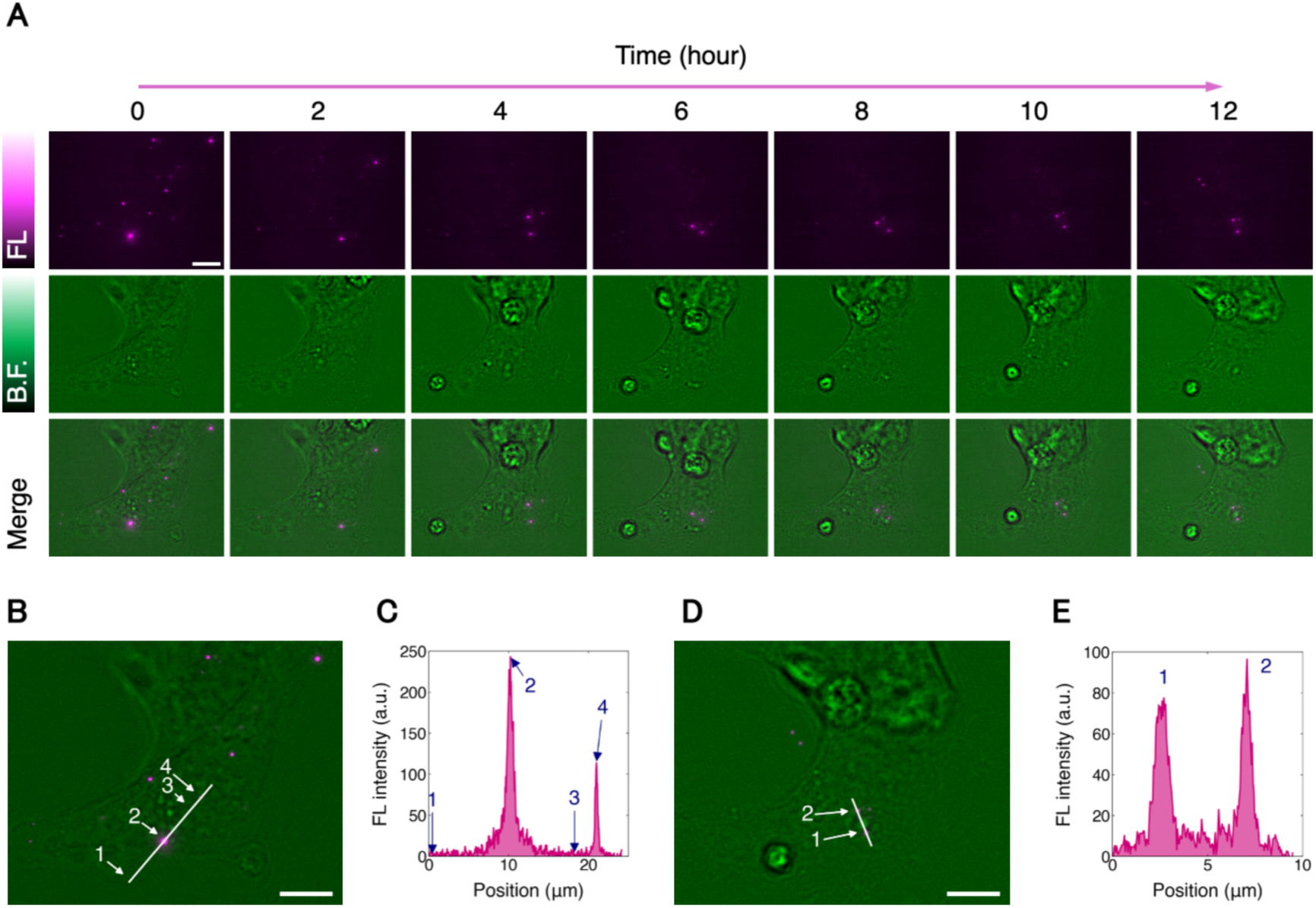
(A) Snapshot images of Adots in living HEK293 cells at 0, 2, 4, 6, 8, 10, and 12 h. Fluorescence images (top panels), bright-field images (middle panels), and merged images (bottom panels) are shown. Imaging was performed under continuous-wave 785 nm excitation at 0.15 kW cm^-2^ with 1-min intervals for 12 h. (B) Merged image at t = 0 h from (A). Regions labeled 1–4 correspond to those analyzed in (C). (C) Intensity profiles along the line indicated in (B). (D) Merged image at t = 12 h from (A). (E) Intensity profiles along the line indicated in (D). Two fluorescent spots were observed, with SNR of 15.6 and 19.7, respectively. All scale bars, 10 μm.

We analyzed the data using a particle-tracking approach to study the dynamics of individual Adots. In each fluorescence image frame, individual fluorescent spots corresponding to individual Adots or small aggregates were detected using a Laplacian-of-Gaussian detector, and their positions were determined from the detected spot coordinates. The centroid localization precision was estimated to be approximately 2.0 nm (see Supporting Information, Section 2). Next, trajectories were reconstructed by linking the detected fluorescent spots across consecutive frames. In this linking process, a cost function was defined based on two criteria: the frame-to-frame displacement of each particle and the change in its fluorescence intensity. Particle trajectories were then optimized by minimizing this cost function (see Supporting Information, Section 3).

Figures 3A–3C show merged bright-field and fluorescence images from the same imaging experiment shown in Fig. 2. Representative frames acquired at 0, 100, and 400 min are shown. Figure 3D shows the trajectories of the two selected particles over the entire imaging period of 12 h. These two particles were separated by nearly 45 µm at t = 0 min, approached to 4 µm by t = 400 min, and then remained within the same domain for the following 320 min. This behavior is consistent with typical intracellular transport of nanoparticles encapsulated in vesicles along the endocytic pathway.^13,22^ The convergence of the two particles suggests that they were transported toward a specific intracellular domain, possibly through active endosomal transport.

**Figure 3.**
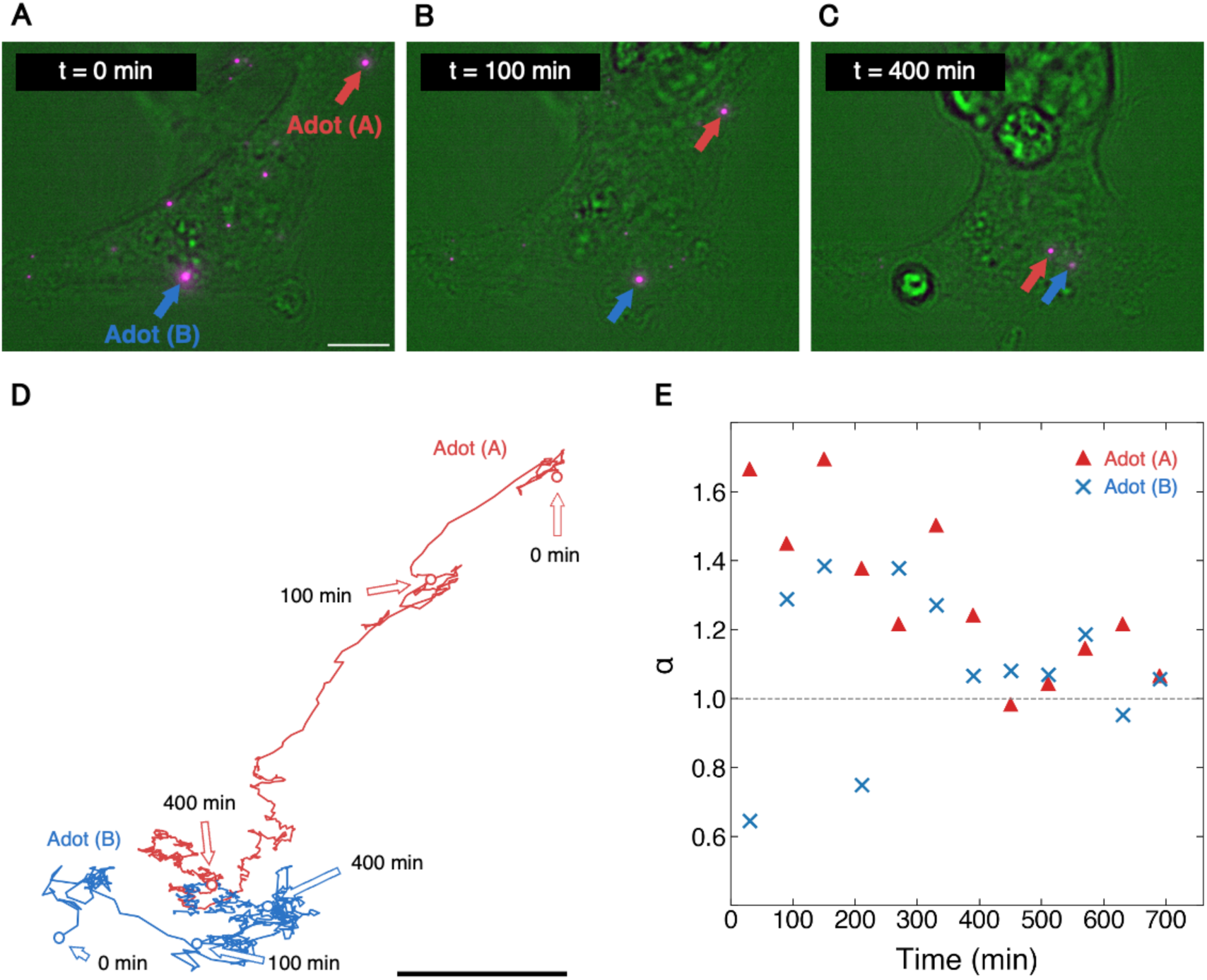
SPT analysis of Adots in live HEK293 cells. (A–C) Representative merged images of Adots in living HEK293 cells acquired at 0, 100, and 400 min. Arrows indicate the particles corresponding to the trajectories shown in (D). (D) Long-term trajectories of two selected particles over the entire 12-h observation period. (E) MSD analysis of the trajectories shown in (D). MSD was calculated for every 60 min interval, and the anomalous diffusion exponent α was extracted by fitting the data to MSD = *Aτ*^!^. All scale bars, 10 μm.

To further characterize the transport behavior, the mean squared displacement (MSD), which represents the average squared displacement by a particle over a given lag time, was calculated for each 60-min interval. The anomalous diffusion exponent α was then extracted by fitting the MSD curves to

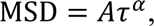

where *A* is a constant and *τ* is the lag time (Fig. 3E). The exponent *α* was used to characterize particle transport behavior at minute-scale sampling intervals over several-hour observations.^1^ For an ideal Brownian random walk, *α* is equal to 1 because the MSD increases linearly with lag time. In contrast, *α* approaches 2 for unidirectional motion at a constant velocity, where the displacement increases linearly with time and the MSD therefore scales quadratically. When *α* is smaller than 1, the particle motion is classified as subdiffusive. Such subdiffusive behavior can arise from hindered or confined transport caused by obstacles, molecular crowding, trapping, or interactions with intracellular structures.

As the MSD curves obtained from the present measurements were well reproduced by a single power law (see Supporting Information, Section 4), the fitted *_α_* provides a well-defined descriptor of the effective long-time transport mode at the scales we observe. For the particle shown in red (Adot (A)), *_α_* was greater than 1.2 during the 0–400 min period, indicating partially persistent, directionally biased transport during this period. Similar superlinear MSD behavior and directionally biased motion have been reported for endocytosed gold nanoparticles undergoing intracellular vesicular transport, where high-motility particles were associated with microtubule-dependent transport during endosomal trafficking.^23^ In contrast, *_α_* was approximately 1.0 during the 400–500 min period for both particles, indicating diffusive behavior. Similar alternation between directed and nondirectional phases has been reported in short-timescale SPT of QD-tagged kinesin motors in living cells.^24^ Although the tracked objects and observation timescales differ, the present results suggest that endocytosed Adots also exhibit multiple effective dynamic modes over much longer observation periods. In contrast to the behavior observed before 500 min, the *_α_* values of the two particles showed good agreement after 500 min. Together with their spatial convergence, this agreement suggests that the particles accumulated within the same intracellular domain and subsequently experienced similar physical constraints imposed by a common organelle or cytoskeletal network, resulting in similar transport dynamics.

## 3. Summary

In summary, we demonstrated the advantages of Au_42_(PET)_32_ as a probe for long-term imaging and SPT in living cells. By using NIR excitation at 785 nm, cellular autofluorescence was suppressed to a negligible level and photodamage was lowered under the present imaging condition. Combined with the high photostability of Au_42_, this approach enabled 12 h SPT in live HEK293 cells. We visualized the intracellular transport of Adots and characterized its transport behavior over hour-scale time windows, observing temporal transitions between directionally biased and diffusive modes, as well as the accumulation of particles within a localized intracellular region. These results demonstrate that Au_42_ provides a platform for long-term and low invasive SPT and offers an experimental framework for investigating intracellular transport dynamics over extended periods. Statistically powered population-level analyses on larger trajectory ensembles, together with co-localization with organelle markers, represent natural next steps. With further optimization of surface modification to improve water solubility and chemical specificity, Au_42_ is expected to serve as an even longer-term SPT probe and functional nanoparticle, thereby further advancing the understanding of intracellular transport mechanisms.

## 4. Materials and Methods

### 4-1. Preparation of Adots

The synthesis of Adots was carried out following the procedure previously reported for the preparation of Raman-active dots.^25^ All operations were performed at room temperature. First, 46.8 µL of a 60-nm polystyrene nanoparticle dispersion (4% w/v, Invitrogen, C37231), 75.0 µL of Pluronic F-127 solution (2% w/v, Invitrogen, P6867), and 178.1 µL of ultrapure water were mixed in a 1.5-mL plastic tube. Next, 135.0 µL of tetrahydrofuran (THF) was added, and the mixture was vortexed for 5 min to swell the polystyrene nanoparticles. After that, 20 µL of a THF solution (1.5 mmol/L) of Au_42_, independently synthesized according to the reported procedure,^26^ was added, followed by vortex mixing for 20 min. Subsequently, 9 mL of ultrapure water was added to the mixture to shrink the nanoparticles again, thereby immobilizing the Au_42_ clusters inside the particles. To remove the solvent, the mixture was centrifuged at 2900 g for 10 min using a 100 kDa MWCO centrifugal filter (Millipore, UFC910096). Then, 5 mL of ultrapure water was added and centrifugation under the same conditions was repeated twice. Finally, approximately 200 µL of the Adots-containing suspension remaining on the filter was collected, and ultrapure water was added to bring the total volume to 1 mL, yielding an Adots suspension. Under these synthesis conditions, assuming that all added Au_42_ clusters are encapsulated inside the polystyrene nanoparticles, the local Au_42_ concentration inside the particles is 15 mmol/L.

### 4-2. Cell cultivation

The human embryonic kidney cell line HEK293 was maintained in Dulbecco’s Modified Eagle Medium (DMEM) supplemented with 10% fetal bovine serum (FBS) and 1% penicillin–streptomycin. Cells were cultured at 37 °C under a 5% CO₂ atmosphere.

### 4-3. Introduction of Adots into cells

One day before the imaging experiments, the cells were seeded onto glass-bottom dishes at a cell density such that it reaches 50–70% confluency on the following day. An aqueous Adots suspension was added to the culture medium 120 min before the imaging experiments. Immediately before starting the imaging, the cells were rinsed with PBS to remove excess probes.

### 4-4. Near-infrared fluorescence imaging systems

We performed long-term NIR imaging using a home-built microscope designed to satisfy three requirements: acquisition of Adot fluorescence images, acquisition of bright-field images of cells, and continuous autofocus control (Fig. 4).

**Figure 4.**
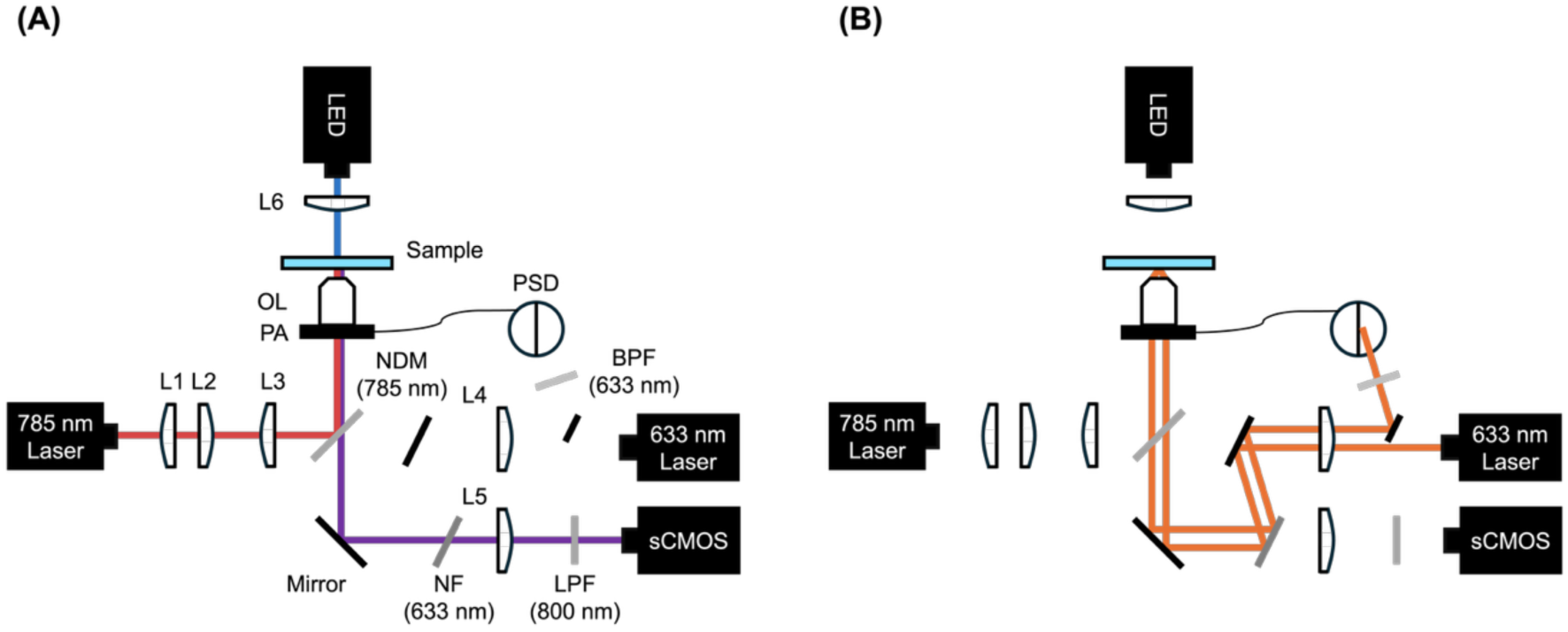
(A) Schematic diagram of our home-built NIR fluorescence microscope. L1-L6: Lenses, NF: Notch filter, NDM: Notch dichroic mirror, PA: Piezoelectric actuator, OL: Objective lens, PSD: Position-sensitive detector, BPF: Band-pass filter, LPF: Long-pass filter, sCMOS: Scientific complementary metal–oxide–semiconductor camera. (B) Schematic diagram of the autofocus system used for continuous focus stabilization.

For fluorescence imaging of Adots, a 785-nm continuous-wave laser was used as the excitation source because Au_42_ exhibits an absorption band centered at 800 nm. Although Au_42_ also has an absorption band at 400 nm, this excitation band was not used because excitation at such short wavelengths can induce higher phototoxicity in living cells. The 785-nm laser beam was expanded using a pair of lenses (L1 and L2) and guided through lens L3 and the objective lens (OL) to achieve wide-field illumination at the sample plane. Fluorescence emitted from Adots was collected by the same objective lens, separated from the excitation light using a notch dichroic mirror (NDM), and detected using an sCMOS camera after passing through the imaging lens (L5). An 800-nm long-pass filter (LPF 800 nm) was placed in front of the camera to further block residual excitation light after the NDM.

For bright-field imaging of cells, an NIR LED was used as the illumination source because NIR light exhibits low cytotoxicity and high transmission through biological samples. The LED light was collimated using lens L6 and directed onto the sample plane. Transmitted light from the sample was collected by the objective lens and imaged onto the camera. The detailed specifications of the optical components are summarized in Table 1.

**Table 1.** Detailed specifications of the optical components used in the microscope system.

| Optical Component | Specification (Model) |
| --- | --- |
| 785 nm Laser | OBIS LX 785 nm 50 mW (Coherent)<br>Emission wavelength 785 nm. |
| LED | M940L3, Thorlabs<br>Center wavelength: 940 nm. |
| 633 nm Laser | 1107P, JDSU<br>Emission wavelength 633 nm. |
| Objective lens (OL) | MRD71670, Nikon<br>Magnification: 60x, NA: 1.42, Working distance: 0.15 mm. |
| Notch dichroic mirror (NDM) | NFD01-785-25x36, Semrock<br>Center wavelength: 785 nm, Reflectance: > 98% at 785 nm. |
| sCMOS Camera (sCMOS) | pco.edge 5.5, pco.<br>Resolution: 2560 × 2160 pixels, Pixel size: 6.5 × 6.5 μm. |
| Position sensing detector (PSD) | PDQ80A, Thorlabs<br>Spectral response range: 400–1050 nm. |

For long-term live-cell imaging over several hours, focus drift caused by changes in the distance between the sample and the objective lens becomes a serious problem. To overcome this limitation, we developed an autofocus system. A 633-nm continuous-wave laser was focused onto the back focal plane of the objective lens near the edge of the back aperture using a lens (L4). At this wavelength, Au_42_ exhibits only weak absorption. The laser beam was reflected at the interface between the cover slip and the immersion oil, and the reflected light was detected using a position-sensitive detector (PSD). When the distance between the sample and the objective lens changed, the position of the reflected beam also shifted, resulting in a change in the PSD signal. This signal was used as feedback to drive the piezoelectric actuator equipped with the objective mount and maintain focus during long-term imaging.

### 4-5. Scanning electron microscopy imaging of Adots

SEM images of Adots were taken by using SU8000 (Hitachi High-Tech) operated at an accelerating voltage of 1.5 kV.

## ASSOCIATED CONTENT

### Supporting Information

The Supporting Information is available free of charge at https://pubs.acs.org/doi/10.1021/acs.analchem.XXXXXXX.

Detection of fluorescent particles during long-term imaging; estimation of the centroid localization error; trajectory analysis using a cost function and confidence score; and mean squared displacement (MSD) analysis (Figures S1–S6) (PDF)

## AUTHOR INFORMATION

### Author Contributions

K.H. conceived and directed the project. S.Y., R.N., R.O. and K.H. designed the study. S.Y. and K.H. performed the experiments. S.Y. prepared the Adots samples. S.T. and T.T. prepared and characterized the nanocluster samples. S.Y. and K.H. analyzed the data. All authors wrote the paper.

## ACKNOWLEDGMENT

The authors thank Y. Murakami, T. Tamura, N. Sagami for their helpful discussion and technical support.

## TOC graphic

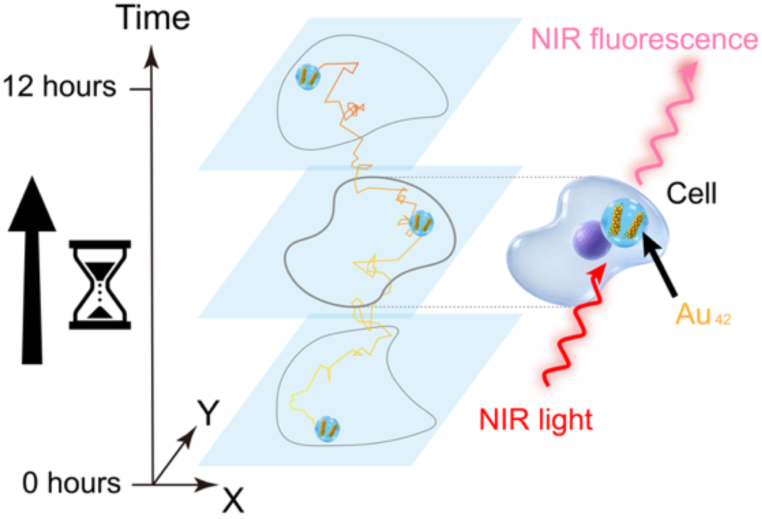

