## Supplementary Information for "Long-term single-particle tracking by NIR imaging using Au_42_ (gold) quantum needles"

### 1. Detection of fluorescent particles during long-term imaging

In each fluorescence image frame, fluorescent spots corresponding to individual Adots or small aggregates were detected using a Laplacian of Gaussian (LoG) detector implemented in the TrackMate plugin of ImageJ. Their positions were determined from the detected spot centers. Figure S1 shows the temporal evolution of the number of detected particles during the 12 h observation period. At least three particles were detected at all time points throughout the measurement.

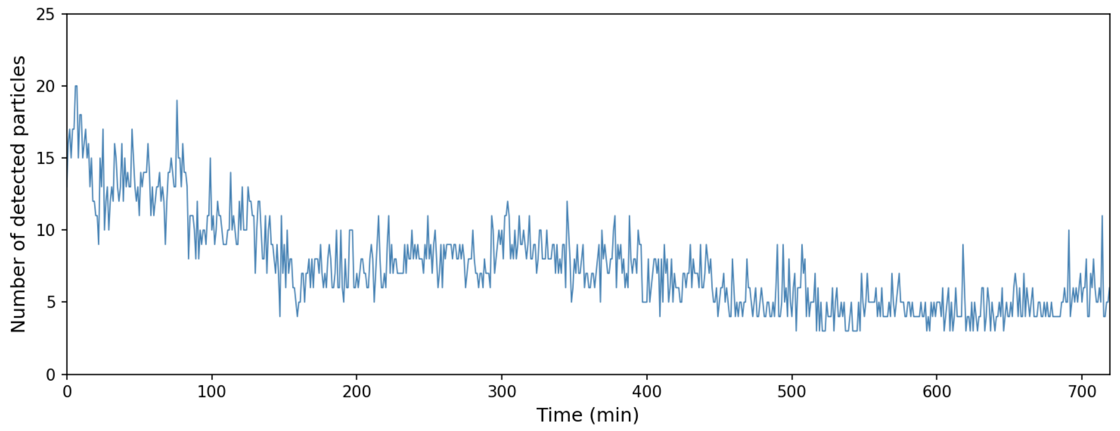


Figure S1. Temporal evolution of the number of detected fluorescent particles during the 12 h observation period.

### 2. Estimation of the error in centroid determination

The localization uncertainty of the fluorescence spots of Adots was estimated using the following equation[1]:

$$\Delta x = \sqrt{\frac{s^2 + a^2/12}{N} + \frac{8\pi s^4 b^2}{a^2 N^2}},$$

where $s$ is the standard deviation of the point spread function (PSF), $a$ is the pixel size, $N$ is the number of detected photons, and $b$ is the background noise per pixel. The PSF was evaluated from fluorescence images of Adots dispersed on a glass substrate acquired with an exposure time of 50 ms. Individual fluorescence spots were fitted with a Gaussian function, yielding $s = 195$ nm. The pixel size at the sample plane was estimated to be $a = 43.3$ nm. The average number of detected photons was estimated to be $N = 17{,}600$ photons from several representative Adots, and the average background intensity was $b = 5.43$ photons/pixel. Based on these values, the centroid localization error $\Delta x$ was estimated to be approximately 2.0 nm.

### 3. Trajectory analysis

Particle trajectories were reconstructed by linking fluorescent spots detected in consecutive fluorescence image frames. The linking procedure was based on a cost function that accounts for both particle displacement and changes in fluorescence intensity. The reliability of the reconstructed trajectories was then quantitatively assessed using a confidence score.

The modeled situation is illustrated in Fig. S2A. At time $t$, two particles ($i = 1, 2$) are present, and at the subsequent time point $t + 1$, two particles $(j = 1, 2)$ are observed. In this case, there are two possible trajectory assignments, as illustrated in Fig. S2B. One assignment corresponds to the combination of solid arrows, whereas the other corresponds to the combination of dashed arrows.

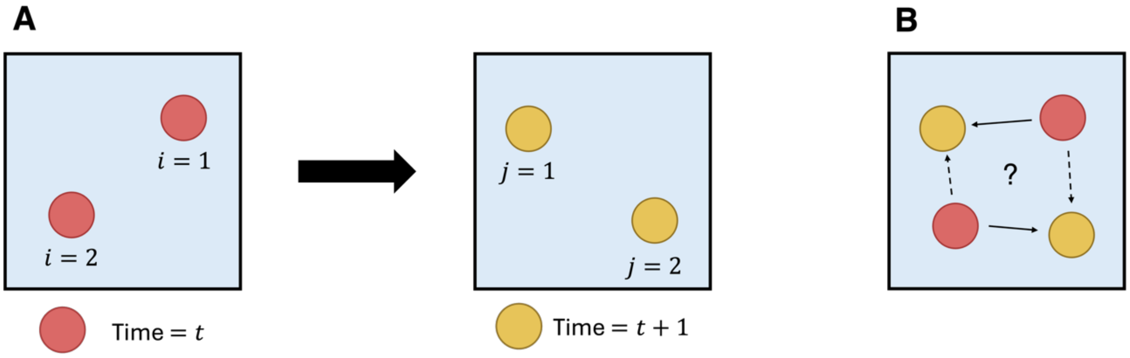


Figure S2. (A) Schematic illustration of the modeled situation. Particles observed within the field of view at time $t$ are shown in red, whereas those at time $t + 1$ are shown in yellow. (B) Schematic illustration of the tracking problem considered in this study. For each particle at time $t$, the corresponding particle at time $t + 1$ was determined.

To quantitatively evaluate which combination of trajectories was most plausible, the following cost function was defined for each candidate link:

$$C(i,j;t) = w_d \frac{d(i,j;t)}{d_{\max}} + w_r \frac{r(i,j;t) - 1}{R_{\max} - 1}, \tag{S1}$$

where $d(i,j;t)$ is the distance between the particle $i$ and $j$, $r(i,j;t)$ is the fluorescence intensity ratio between the particle $i$ and $j$, $w_d$ and $w_r$ are weighting parameters for the distance and intensity terms, respectively, $d_{\max}$ and $R_{\max}$ are model parameters. As the distance $d(i,j;t)$ increases, the first term becomes larger. Particles separated by a distance greater than $d_{\max}$ are never linked (Fig. S3A). Similarly, as the fluorescence intensity ratio $r(i,j;t)$ increases, the second term becomes larger (Fig. S3B). The intensity ratio was defined to be greater than or equal to 1:

$$r(i,j;t) = \frac{\max\{I(i;t), I(j;t+1)\}}{\min\{I(i;t), I(j;t+1)\}}, \tag{S2}$$

where $I_{(i;t)}$ is the fluorescence intensity of particle $i$ at time $t$. Candidate links with intensity ratios of larger than $R_{\max}$ were excluded. For each frame pair, the cost function was calculated for all possible links, where the assignments with the minimum total cost was selected. To compensate for the temporary disappearance of particles due to defocusing or fluorescence intensity fluctuations, an additional gap-filling procedure was implemented.

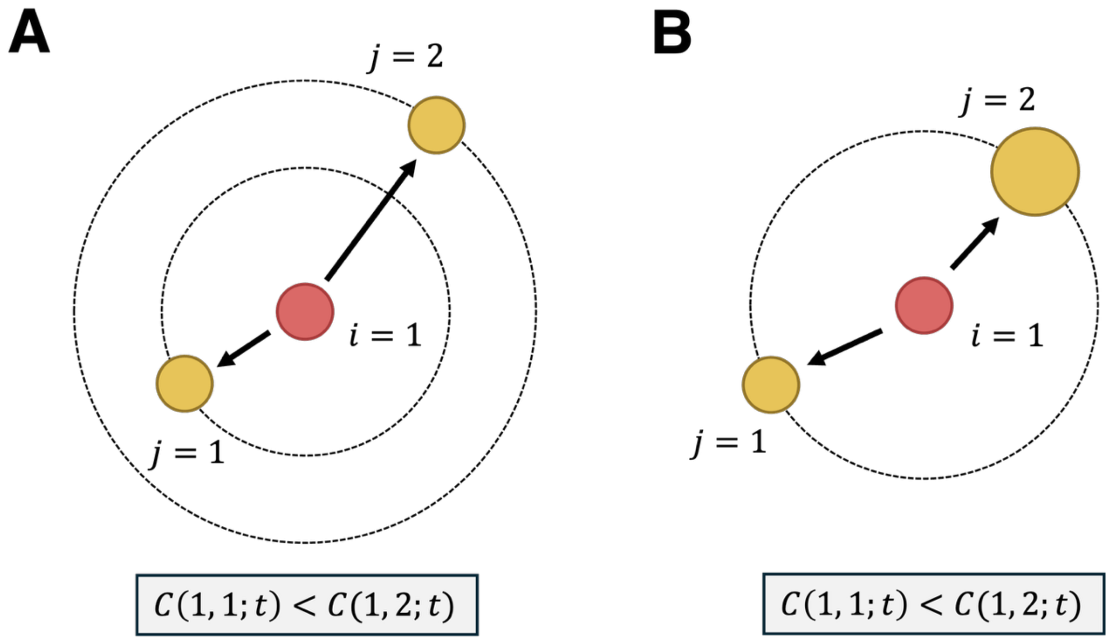

Figure S3. Properties of the cost function used for particle linking. (A) The cost function increases as the distance between particles increases. (B) The cost function increases as the difference in particle intensity increases. The particle size in the schematic represents the particle intensity.

Next, the robustness of the reconstructed trajectories was evaluated using the following confidence score $S(i;t)$:

$$S(i;t) = 1 - \frac{C(i;t)_{\text{best}}}{C(i;t)_{\text{2nd}}}, \tag{S3}$$

where $C(i;t)_{\text{best}}$ and $C(i;t)_{\text{2nd}}$ represent the smallest and second-smallest values of $C(i,j;t)$ among all candidate assignments, respectively (Fig. S4A). A confidence value close to 1 indicates a unique assignment (Fig. S4B), whereas a value close to 0 indicates ambiguity (Fig. S4C).

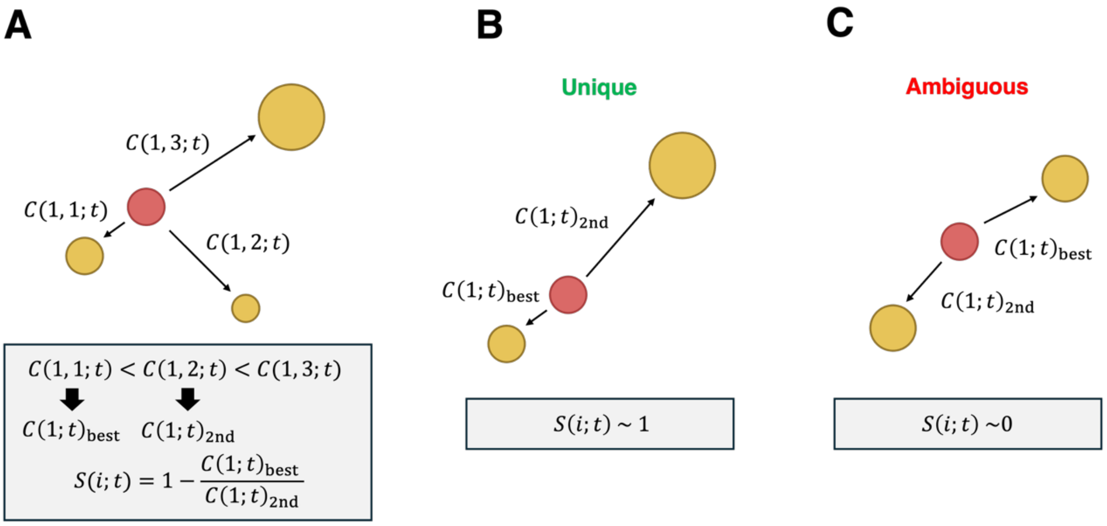


Figure S4. (A) Definition of confidence score $S(i;t)$. The confidence score was defined from the ratio between the smallest and second smallest cost functions among the candidate particle links. (B) A confidence score close to 1 indicates high reliability of the particle trajectory. (C) A confidence score close to 0 indicates low reliability of the particle trajectory.

The confidence scores for the two trajectories shown in Fig. 3D in the main text were calculated and are presented in Fig. S5. In almost all frame pairs, the confidence score was close to 1, indicating that the reconstructed trajectories were reliable. Frame pairs with low confidence scores were mainly observed when the particle density was high or when particle defocusing caused substantial changes in fluorescence intensity.

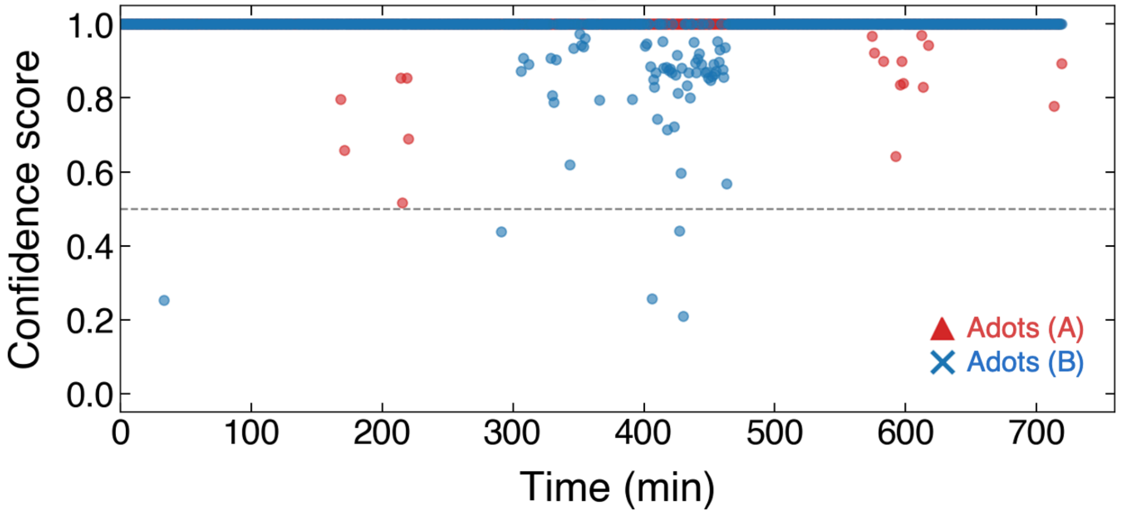


Figure S5. Confidence score for the two trajectories shown in Fig. 3D.

**4. MSD analysis**

The trajectories shown in Fig. 3D in the main text were divided into twelve 60-min segments for calculating the MSD within each time window. The resulting MSD curves were fitted using the following equation:

$$\mathrm{MSD} = A\tau^{\alpha}, \quad \text{(S4)}$$

where $\tau$ is the lag time, $A$ is a constant, and $\alpha$ is the anomalous diffusion exponent. For each time window, the value of $\alpha$ was determined by fitting the MSD curve to Eq. (S4). For large values of $\tau$, the MSD is calculated from a smaller number of data points, which reduces the statistical accuracy. Therefore, the fitting was performed only over time lags corresponding to up to one-tenth of the total duration of each trajectory.

The MSD curves of the two trajectories shown in Fig. 3D were plotted on a log-log scale (Fig. S6). The fitting curves obtained from Eq. (S4) are also shown in the figure.

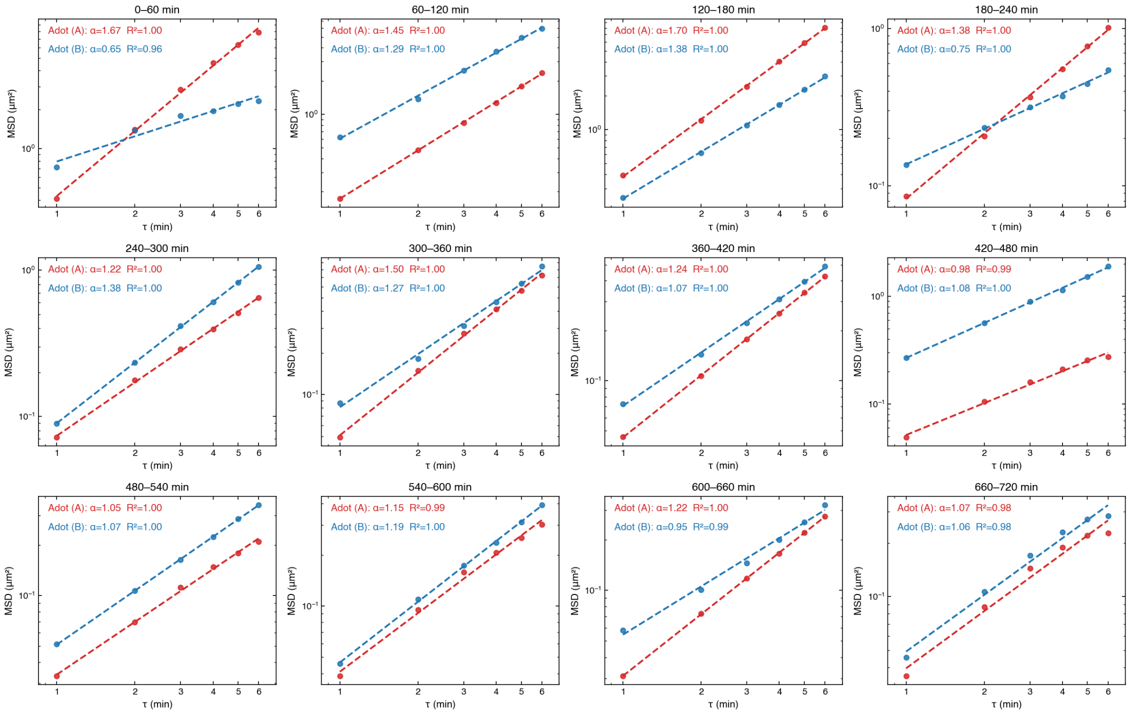

Figure S6. MSD curves of the two trajectories shown in Fig. 3D. The trajectories were divided into 60-min time windows, and the MSD within each window was plotted on a log-log scale. Dashed curves represent the fitting results obtained using Equation (S4).
